# TO BE OR NOT TO BE A PI: ADVICE FOR TRAINEES IN THE BIOMEDICAL/BASIC SCIENCES

**DOI:** 10.64898/2025.12.05.692199

**Authors:** Hei-Yong G. Lo, Lily L. Nguyen, Kristine M. Sikora, Bruce Mandt, Angeles B. Ribera

**Affiliations:** Medical Scientist Training Program, School of Medicine, University of Colorado; Graduate School, University of Colorado, Anschutz Medical Campus; Department of Pharmacology, School of Medicine, University of Colorado; Office of Research Education, School of Medicine, University of Colorado; Department of Physiology & Biophysics, School of Medicine, University of Colorado

## Abstract

When a biomedical or basic science doctoral (PhD) student considers whether or not to become a Principal Investigator (PI), they typically base their decisions on anecdotal data from a limited number of faculty mentors. Alternatively, they may look online for guidance and be met with biased opinions rather than unbiased, comprehensive data. We sought to address this gap by anonymously surveying active faculty at the University of Colorado Anschutz Medical Campus (CU Anschutz) to gain more clarity on the daily life of a PI, the decisions they made as they navigated their career path, and which skills were valuable for securing a faculty position. We also identified important factors in choosing a postdoctoral lab. We found that the majority of PIs would recommend becoming a PI and thought positively of the career path, with most feeling content. But attitudes differed between faculty of different ranks. Further, in addition to scientific vision, hard work, and publications, luck was also deemed as an important factor in becoming and succeeding as a PI. We present our findings so that they may serve as a resource for graduate trainees who are considering a PI/academic research career path. In addition, the information may guide future training plans to optimize achievement of their dream career.

## INTRODUCTION

Becoming an academic principal investigator (PI) in the biomedical/basic sciences has traditionally followed an apprenticeship model where the bulk of the graduate student’s tutorship comes from their direct mentor. As a result, PhD candidates and postdoctoral scholars (trainees) rely on a limited pool of resources to determine whether to pursue a career as an academic PI. These resources may include the experiences of their direct mentor, adjacent mentors or collaborators, and workshop panels featuring individuals from different careers (when available). These traditional resources are subject to their unique biases. For example, a mentor may provide different feedback to a trainee depending on their mentor/mentee relationship (Tuma et al. 2021). Online resources are not immune to biases. Many resources are written as opinion pieces (Luppi et al. 2021; Bourne and Friedberg 2006; Tregoning and McDermott 2020; Odom 2014; Saez et al. 2020). These, while useful and full of wisdom, may not capture the most up to date and accurate information in the quickly-evolving research training landscape. Additionally, they often do not provide data from a broad range of PIs. Taken together, these issues make it challenging for a trainee to arrive at an informed decision of whether or not to become a PI conducting biomedical/basic science research.

One solution to this issue is more exposure. Increased exposure to more faculty has been shown to reduce these biases and increase career preparedness of graduate and postdoctoral scholars (Layton et al. 2020; Fuhrmann 2016). Recent published work aggregating responses from a greater number of faculty has helped overcome individual biases and provide greater transparency. To help trainees prepare for a faculty position, some have evaluated what factors are predictive of success as an applicant in the PI hiring process (Wu et al. 2023; Boysen et al. 2019), and others have measured success during PI rank (Dijk et al. 2014). While incredibly informative, these studies do not limit their focus to PIs and may be field specific.

We sought to provide more robust data to help graduate students and postdoctoral scholars arrive at a decision. To obtain information from the relevant source, we conducted a survey querying PIs about their experiences and sought their advice. The resultant aggregate, quantifiable data provide biomedical/basic science trainees with a more concrete and broad understanding of what led others to become and succeed as a PI conducting biomedical/basic sciences research. By analyzing and sharing the survey results, trainees will have information allowing a more informed decision on whether to become an academic PI. Additionally, this study collectively provides a more comprehensive picture of the skills and factors that PIs value and help trainees develop a more concrete, focused training plan and faculty application.

## METHODS

### Survey design and distribution

We used a non-probability sampling approach at a single institution due to the feasibility challenges inherent in reaching PIs at numerous institutions. We solicited 409 current faculty members who perform biomedical/basic science research at the University of Colorado Anschutz Medical campus (CU Anschutz) in Aurora, Colorado between April to June of 2024. Participants were recruited using emails, posters, and verbal invitations. Surveys were generated and administered using Qualtrics survey software. We received 134 PIs survey responses yielding a response rate of 33%. To ensure all participants were from our target population, we asked each PI to confirm that they are PIs as defined as having a faculty appointment at CU Anschutz and conducting biomedical or basic science research.

The initial draft of the survey, written by HYG Lo, was revised following feedback from a total of 6 PIs who perform(ed) biomedical/basic science research (see acknowledgements). The 6 PIs provided feedback about the comprehensiveness of the survey questions, how well they addressed the overarching goal of the survey, clarity of the questions, and any other additional comments. The surveyed items reflect topics and questions often addressed in workshops held at conferences and retreats. Questions that could be quantified were prioritized over free-responses and the total number of questions (26) was kept minimal to prevent survey fatigue. Most questions were not mandatory, allowing participants to skip certain questions. For the survey to be considered “completed,” the respondent needed to progress through each page. Because not every item in the survey was mandatory, some respondents completed the survey, but did not answer every item in the survey. We used every response to every item recorded as long as the respondent met criteria for having completed the survey.

The survey took approximately 10 minutes to complete. For access to the complete survey, please contact corresponding author Hei-Yong Lo.

Conditions were set to allow PIs to respond anonymously and prioritize frank and honest responses. For example, the survey did not collect names, department affiliation, IP addresses, or other demographics that would easily identify an individual. The survey limited demographic information that would allow for stratification of the data unique to those demographics (e.g. academic rank, gender or minority status).

PIs were also asked some open-ended questions regarding their career as a PI, and some PIs chose to allow their responses to be published (**Supplemental File 2**). The survey was granted IRB exemption by the CU Anschutz IRB office under COMIRB 24-0319.

### Analysis

All analyses were performed with RStudio. Original data were curated to correct for inputs that were not compatible with analyses (e.g. 40-50 was converted to 45; 45% was converted to 45). For responses where an obvious conversion to an integer could not be made, the response to that particular question was dropped. This only affected three questions.

Statistical analyses were performed using RStudio statistical packages. Multi-way statistical analyses were performed with two-way ANOVA tests. T-tests were used for pairwise comparisons with multiple comparisons made by applying the Benjamin-Hochberg procedure. For Likert scale data, which is ordinal, we used the Wilcoxon rank sum test with continuity correction to account for non-parametric data (Sullivan and Artino 2013; Jamieson 2004). Where appropriate, the mean is displayed in addition to the median.

## RESULTS

### Description of participants

Over the span of two months in 2024, we reached out to 409 principal investigators at CU Anschutz through emails, face-to-face solicitations, and flyers. Of the solicited PIs, 134 participants completed the survey (see Methods). At the time of this survey, CU Anschutz was and continues to be designated as an R1 research institution.

Some studies have noted that PI demographics may impact their attitudes. For example, being a woman in academia has been correlated with negative responses about work/life balance, mental and physical health (Ysseldyk et al. 2019; Martinez et al. 2007). Accordingly, we asked respondents to self-identify their gender and minority status, for stratification of data in subsequent analyses. The full breakdown of self-ascribed demographics can be found in **Table 1**. While men are more prevalent than women in the biomedical sciences field nationally (Martinez et al. 2007), we received an equal proportion of male (n=65) and female (n=67) responses (2 participants did not share this information). This is representative of the faculty at CU Anschutz, based on internal data. Close to 10% of our respondents self-identified as minority status as per guidelines formerly provided by NIH (n=13), which is lower than the North American populational average (Getz et al. 2022). To test whether the position and rank correlated with different responses (Burns et al. 2018), respondents provided their academic rank; we received meaningful representation from each level of professorship (28 assistant professors, 45 associate professors, and 61 full professors), as well as time spent as a PI. Finally, because we queried PIs at an academic medical center, we asked respondents to indicate whether they had clinical responsibilities to test whether that factor correlated with different responses. The 23 respondents (17% of completed surveys) are likely MD or MD/PhDs.

**Table 1:**
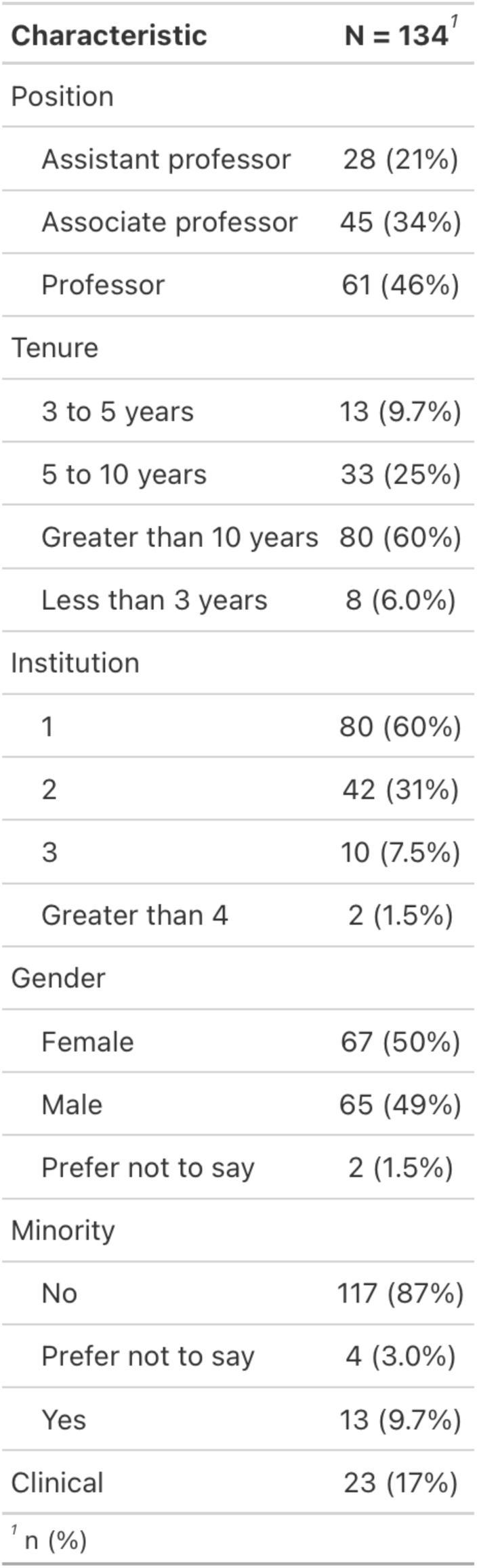
Responders’ characteristics. Table depicting the total number PI respondents who finished the survey (134). The number of PIs with self-ascribed demographics are shown.

| **Characteristic** | **N = 134<sup>†</sup>** |
| --- | --- |
| Position |  |
| Assistant professor | 28 (21%) |
| Associate professor | 45 (34%) |
| Professor | 61 (46%) |
| Tenure |  |
| 3 to 5 years | 13 (9.7%) |
| 5 to 10 years | 33 (25%) |
| Greater than 10 years | 80 (60%) |
| Less than 3 years | 8 (6.0%) |
| Institution |  |
| 1 | 80 (60%) |
| 2 | 42 (31%) |
| 3 | 10 (7.5%) |
| Greater than 4 | 2 (1.5%) |
| Gender |  |
| Female | 67 (50%) |
| Male | 65 (49%) |
| Prefer not to say | 2 (1.5%) |
| Minority |  |
| No | 117 (87%) |
| Prefer not to say | 4 (3.0%) |
| Yes | 13 (9.7%) |
| Clinical | 23 (17%) |
| <sup>†</sup> n (%) |  |

### Most PIs would recommend their career choice

Half of the respondents (n=71, 50.0%) recommended becoming a PI (**Figure 1A**). Only 5 respondents (3.8%) would not recommend becoming an academic PI. The rest of the respondents either did not complete the question (n=3) or believed it would depend (n=55, 42%). As a group, full professors were more likely than assistant professors (p adj = 0.047) or associate professors (p adj = 0.018) to recommend becoming a PI (**Figure 1A**). Whether or not a PI would recommend an academic career path was not statistically different between genders, minority status, academic rank, or whether PIs had clinical responsibilities.

**Figure 1:**
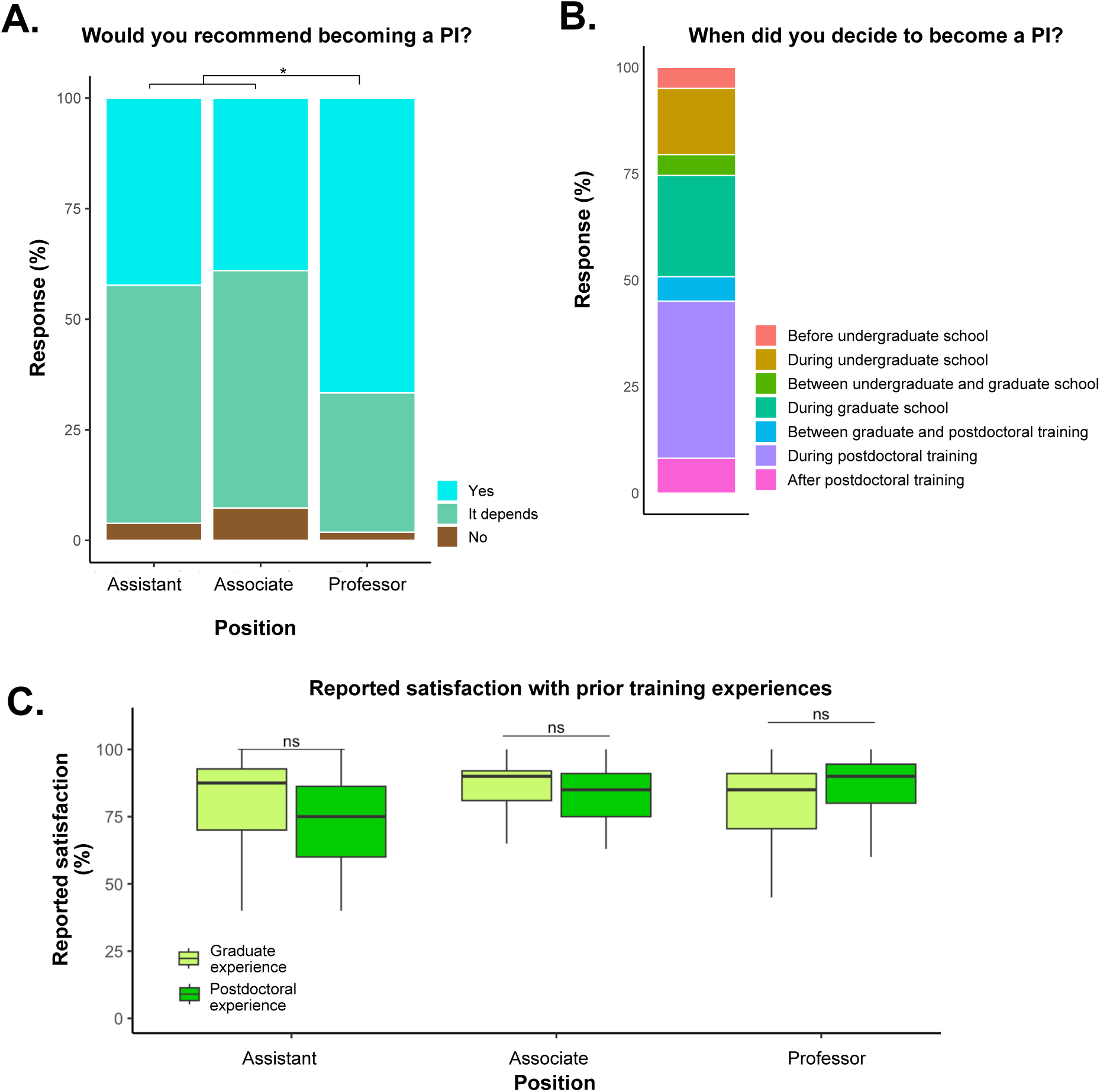
Would PIs recommend becoming a PI. A. Bar graphs depicting the proportion of PIs who would recommend becoming a PI stratified by PI rank. B. A bar graph depicting when along their training PIs made the decision to become a PI. Percentages reflect the percent of PIs who selected that period of their training as the definitive moment they made the decision. C. Boxplots of how PIs rated their graduate and postdoctoral experience (out of a satisfaction scale of 0-100). Responses were stratified by PI rank. Student T-tests with BH correction were used to compare conditions (ns represents p > 0.05, * p <0.05, ** p < 0.01, *** p < 0.001).

### Graduate and postdoctoral experiences inform becoming a PI

To our knowledge, there is no publicly available data on when PIs decided to become a PI. A quarter of PIs decided to become a PI prior to graduate school (n=33, 25%) (**Figure 1B**). Twenty-four percent (n=32) of PIs decided during their graduate training (**Figure 1B**). Half of PIs (n=67, 50%) made their decision after graduate school, with the majority of those deciding during their postdoctoral experience (n=48, 36%) (**Figure 1B**).

PIs rated their satisfaction with their graduate and postdoctoral experiences on a scale of 1 to 100. We found that on average, PIs rated both experiences highly, regardless of academic rank (**Figure 1C**). The PI’s rated their graduate experience as 80/100 (95% CI 76.5 to 85.5) and postdoctoral experience received an average rating of 81/100 (95% CI 77.1 to 84.0) (**Figure 1C**). The data revealed no significant differences between satisfaction ratings of graduate school versus postdoctoral experiences (**Figure 1C**).

### The majority of PIs are satisfied with their occupation

To understand general attitudes and concerns of PIs at an R1 research institution, our survey included a short climate survey. The majority of PIs (85%) agreed (either somewhat or strongly) with the statement that they feel fulfilled with their career (**Figure 2A**). Conversely, only 9% of PIs either somewhat or strongly agreed with the statement that they regret becoming a PI (**Figure 2A**). Most felt they were confident they would continue to be a PI in the next 5 years (80% agree, 14% disagree). The majority of PIs felt a sense of financial stability (60% agree, 29% disagree), job security (63% agree, 28% disagree), and control over their career (66% agree, 19% disagree). Around half of PIs felt they had good work life balance (53% agree, 32% disagree). Still, despite these positive attitudes, it is important to note that a quarter of PIs disagreed with job security and having a good work-life balance. A quarter (25%) of PIs agreed that they thought about leaving the career path often (**Figure 2A**). While some also disagreed with having financial stability, this question is hard to interpret because the question did not specifically ask whether this was in their salary or specifically related to research funding.

**Figure 2:**
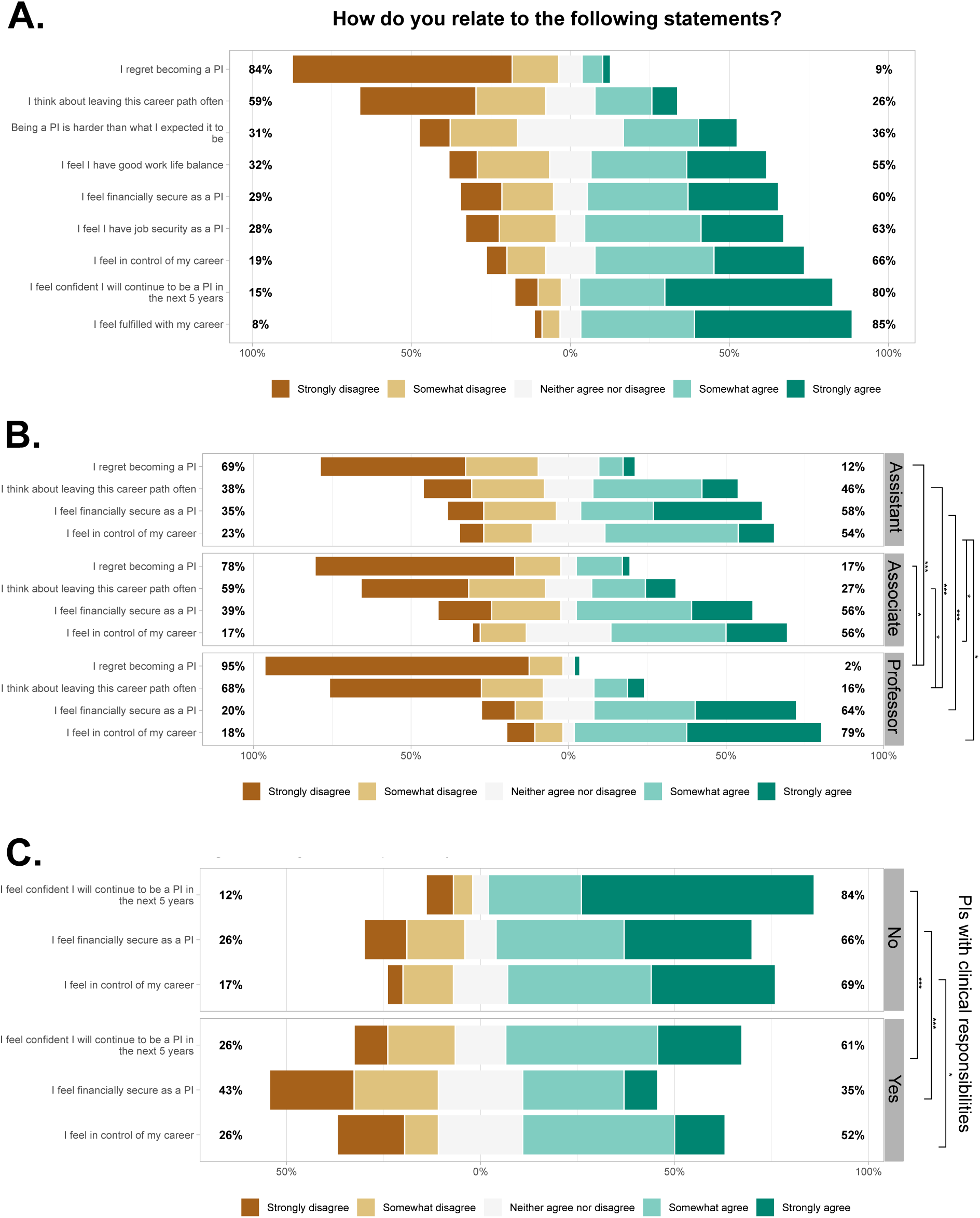
PIs are generally content with their occupation. A. Likert scale chart of queried questions across all PIs. Percentages reflect the aggregate percentage of PIs who “somewhat” or “strongly” agreed or disagreed with an item. B. Likert scale of specific questions whose responses were found to be statistically significant. Results stratified by rank. C. Likert scale of specific questions whose responses were found to be statistically significant. Results stratified by PIs with or without clinical responsibilities. Wilcoxon rank sum test with continuity correction were used to compare conditions (ns represents p > 0.05, * p <0.05, ** p < 0.01, *** p < 0.001).

PIs were equally divided in whether or not becoming a PI was harder than their expectations. Thirty-three percent of PIs disagreed that becoming a PI was harder than their expectations, whereas 35% agreed that becoming a PI was harder than expected (**Figure 2A**).

We queried whether certain factors changed how satisfied PIs were with their profession. We stratified the same items in the question by gender, minority status, clinical responsibilities, and academic position (**Supplemental File 1**). We found no statistical differences between gender or minority status for any item on this question and the majority of PIs were content with being a PI. However, we noted significant differences by academic rank. Compared to full professors, assistant and associate professors were more likely to “regret becoming a PI” (p=0.00022 and p=0.015, respectively) and were more likely to “think about leaving this career path often” (p=0.0078 and p=0.023, respectively) (**Figure 2B**, **Supplemental File 1**). Assistant professors were also less likely to feel in control of their careers (p=0.012) (Figure 2B, **Supplemental File 1**). Because differences were noted at academic rank, we wondered whether length of employment at a certain rank (assistant or associate professor) affected their responses. Assistant professors who had worked > 5 years were less likely to “think about leaving this career path often” (p=0.033) than those who had worked < 5 years (**Figure S1A**, **Supplemental File 1**). Otherwise, no differences were noted between groups of the same rank.

We found that those with clinical responsibilities were less likely to feel financially secure (p=0.0069), less likely to feel in control of their career (p=0.035) and less likely to feel confident they would continue to be a PI in the next 5 years (p=0.00059) (**Figure 2C**, **Supplemental File 1**). However, it is important to note that financial security was not explicitly defined in this survey, and it is not clear if financial concerns were professional vs related to research funding or both.

### Breaking down how PIs spend their time

There is little information regarding how many hours PIs work and how that time is divided. To address this, we asked how many hours are spent performing activities related to being a PI. On average, PIs recorded spending 46.14 hours a week (95% CI 43.35h to 48.92h) fulfilling PI responsibilities (**Figure 3A**). There were no statistical differences between academic rank, gender, or minority status (**Figure S2A-C**). However, PIs with clinical responsibilities spent significantly less time performing PI duties than those without clinical responsibilities (**Figure S2D**). It is important to note that the number of hours spent on “duties related to PI” was self-defined by the PI and otherwise unspecified (e.g. some PIs may be solely limiting to time spent in the lab).

**Figure 3:**
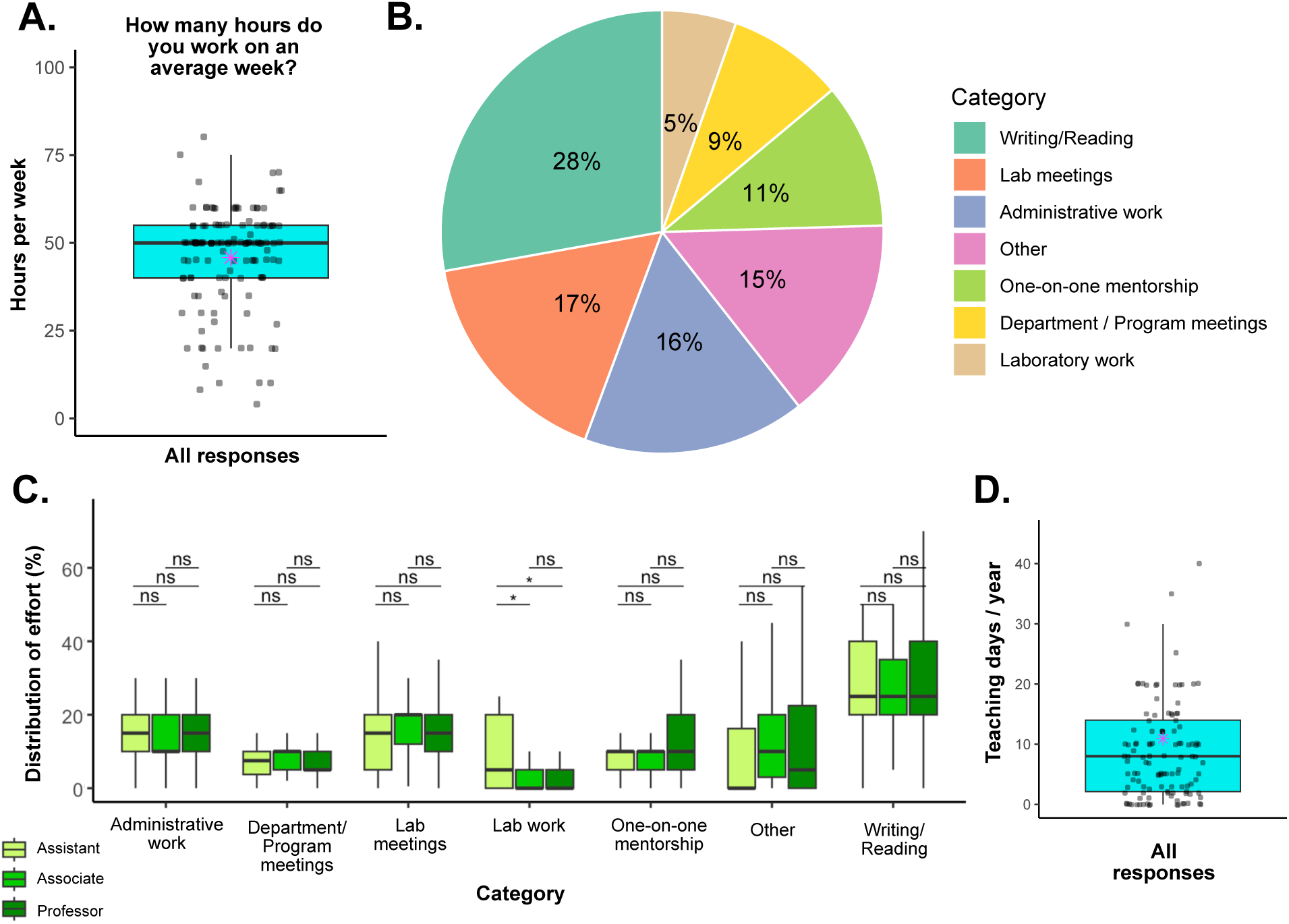
Breakdown of PI responsibilities. A. A boxplot depicting how many hours each PI (black dot) reported spending performing PI duties per week. Mean shown in magenta. B. A pie graph depicting the reported effort (as a percentage) the average PI spent performing each activity. C. Boxplots depicting the reported effort (as a percentage) PIs spent performing each activity in B stratified by academic rank. Student T-tests with BH correction were used to compare conditions (ns represents p > 0.05, * p <0.05, ** p < 0.01, *** p < 0.001). D. Boxplot depicting how many days each PI (black dot) reported teaching per year. Mean shown in magenta.

We next asked how PIs distribute this time across various tasks on a given day. The tasks queried were as follows: administrative work, writing/reading, laboratory work, department / program meetings, lab meetings, one-on-one mentorship, and “other” tasks. We found that on an average day, the average PI spent more of their time reading/writing (28%, 95% CI 25.47% to 30.26%) than on any other individual task (**Figure 3B**). On average, PIs spent a total of 25% of their time on meetings (8% departmental (95% CI 7.33% to 9.56%), and 17% lab meetings (95% CI 15.05% to 18.41%) (**Figure 3B**). They spent an average of 16% (95% CI 14.44% to 18.12%) doing administrative tasks and 15% (95% CI 11.05% to 18.27%) on other tasks not covered in this list (**Figure 3B**). Eleven percent of their time was spent on one-on-one mentorship (95% CI 9.39% to 12%) (**Figure 3B**). PIs spent the smallest proportion of their time conducting physical laboratory work (5.52%, 95% CI to 3.72% to 7.32%) (**Figure 3B**). For most items, we found that how PIs spent their time did not statistically differ across academic rank except for time spent doing laboratory work (**Figure 3C**). Assistant professors were significantly more likely to have recorded more time spent performing laboratory work compared to associate professors or full professors (**Figure 3C**). Those with clinical responsibilities reported spending significantly more time on one-on-one mentorship (**Figure S2E**). No significant differences were noted between genders or minority status (**Figure S2F-G**).

PIs often teach. We therefore asked how many days PIs spend teaching. On average, PIs spent an average of 11 days of the year teaching (95% CI 8.68 days to 13.30 days) (**Figure 3D**). The number of days spent teaching did not statistically differ based on academic rank (**Figure S2H**). Note that CU Anschutz is an academic medical center with graduate-level courses without undergraduates that are taught by professors of each rank.

### Impact of being a PI on faculty rank progression

Using the respondents’ self-assigned demographics, within each academic rank with length of employment, we queried the time each PI spent at each faculty rank (**Figure 4A, B**). All respondents who had been PIs for fewer than 3 years (n=8) were assistant professors (**Figure 4B**). For PIs who had worked 3-5 years (n=13), 85% were assistant professors and 15% were associate professors (**Figure 4B**). For PIs who had worked 5-10 years (n=33), 70% were associate professors, 27% were assistant professors, and 3% were full professors (**Figure 4B**). For respondents who had worked greater than 10 years as a PI (n=80), 75% were full professors and 25% were associate professors. None were assistant professors (**Figure 4B**).

**Figure 4:**
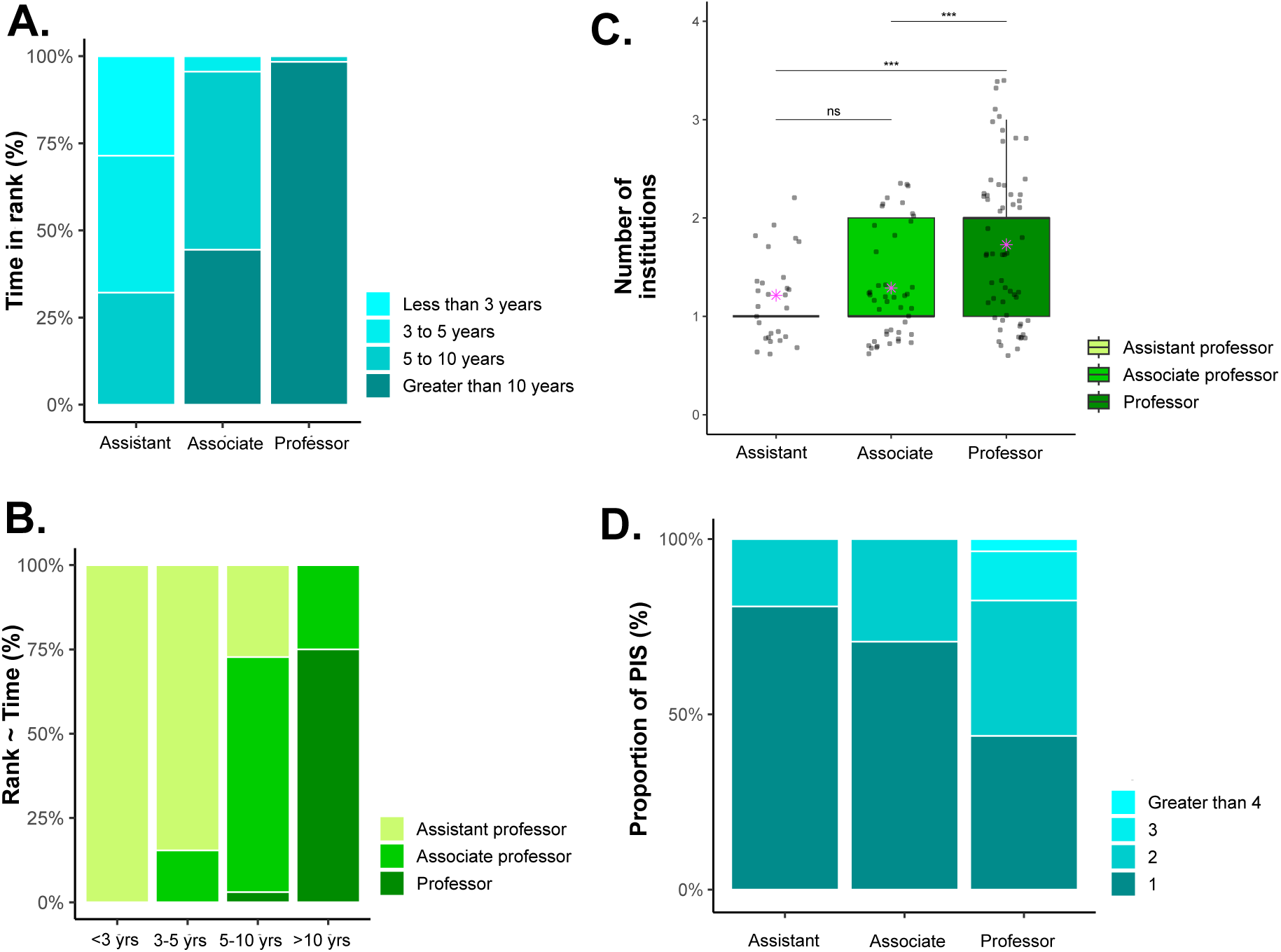
PI career progression as a function of time. A. Bar graphs depicting the percentage of PIs (stratified by academic rank) across time spent as a PI. B. Bar graphs depicting the percentage of PIs (stratified by amount of time spent as a PI) who belong to a specific academic rank. C. Boxplots depicting the number of institutions that PIs (black dot) have reported been employed as a PI at. Note: responses are quantized (1, 2, etc), but black dots are distributed for visualization. Results stratified by academic rank. Mean shown in magenta. Student T-tests with BH correction were used to compare conditions (ns represents p > 0.05, * p <0.05, ** p < 0.01, *** p < 0.001). D. Bar graphs depicting the percentage of PIs (stratified by academic rank) who have worked at a categorical number of institutions.

On average, the majority of assistant professors had been PIs at one institution (n=22, 79%) (**Figure 4C, D**). Similarly, 43% of full professors had only been a PI at one institution (**Figure 4C, D**). Thirty-eight percent of full professors had been a PI at two institutions and 19% had been at three or more institutions (**Figure 4C, D**).

### Factors that PIs considered when choosing a postdoctoral position

We asked PIs how certain factors weighed into their choice of a postdoctoral position. The full responses are shown in **Figure 5A**. Of all respondents (n=134), more than 75% agreed on five items that informed their postdoctoral choice (**Figure 5A**). The top five factors important in choosing a postdoctoral lab reported by PIs were the research of the postdoctoral mentor (95% agree, 1% disagree), the reputation of the mentor (88% agree, 2% disagree), intellectual freedom (84% agree, 5% disagree), location of the postdoctoral experience (80% agree, 11% disagree), and the mentor’s publication record (79% agree, 8% disagree). Conversely, there were a number of items where respondents were almost equally divided about whether that item factored into their choice of postdoctoral position. These factors were salary (8% agree, 65% disagree), promotional/development opportunities (28% agree, 34% disagree), a PI’s graduation record (28% agree, 32% disagree), and the work/life balance (32% agree, 35% disagree) (**Figure 5A**).

**Figure 5:**
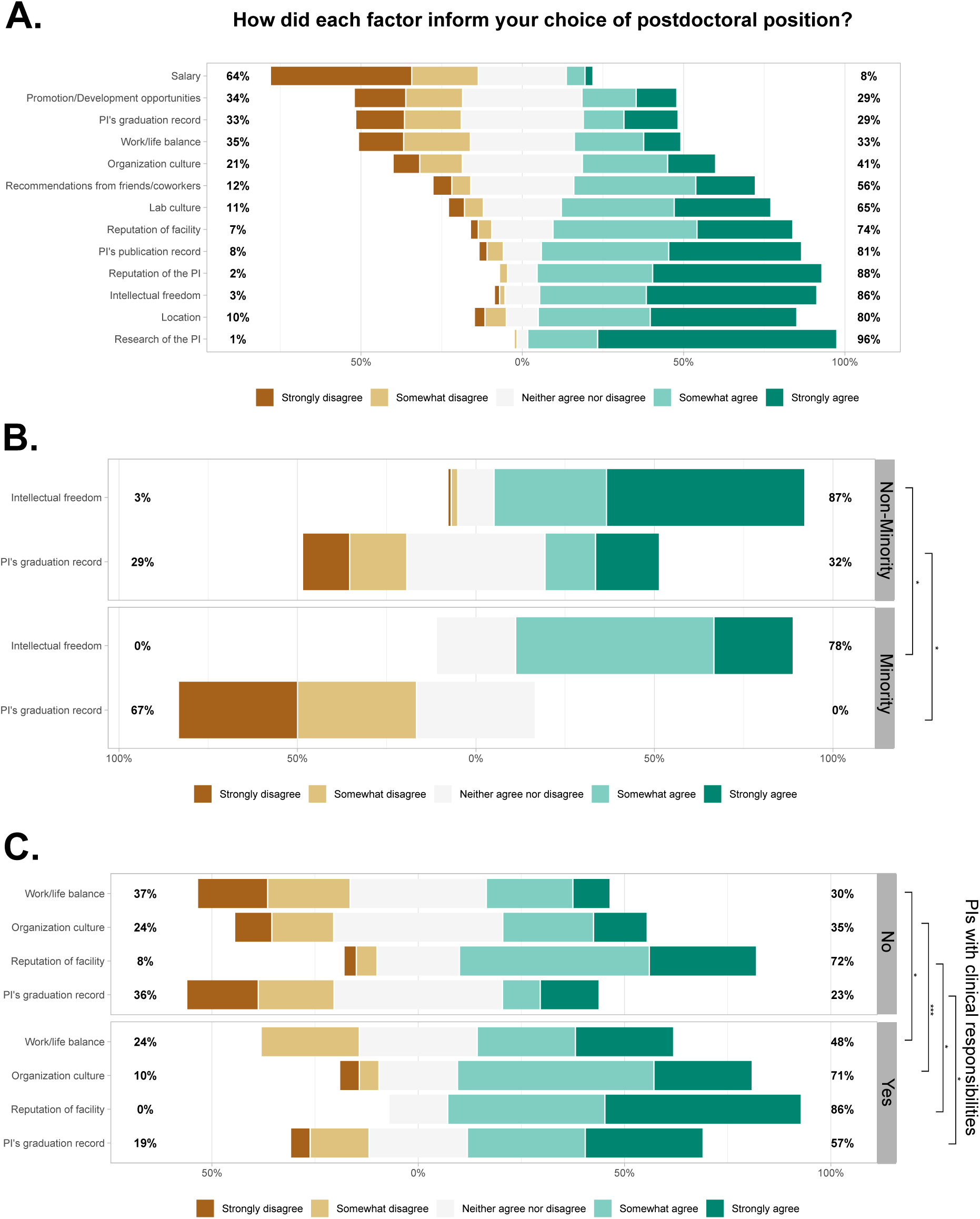
Consideration of factors in postdoctoral lab choice. A. Likert scale chart of queried questions across all PIs. Percentages reflect the aggregate percentage of PIs who “somewhat” or “strongly” agreed or disagreed with an item. B. Likert scale of specific questions whose responses were found to be statistically significant. Results stratified by self-reported minority status. C. Likert scale of specific questions whose responses were found to be statistically significant. Results stratified by PIs with or without clinical responsibilities. Wilcoxon rank sum test with continuity correction were used to compare conditions (ns represents p > 0.05, * p <0.05, ** p < 0.01, *** p < 0.001).

We stratified the same items in the question by gender, minority status, clinical responsibilities, and academic rank (**Supplemental File 1**). We found no statistical significance difference between genders on the items of the question. Respondents who identified as minorities were less likely to factor intellectual freedom (p=0.023) and a PI’s graduation record (p=0.008) than non-minority respondents (**Figure 5B**) in their choice of postdoctoral position. Relative to PIs without clinical responsibilities, PIs with clinical responsibilities were more likely to factor in work life balance (p=0.044), the culture of the organization (p=0.0074), the reputation of the facility (p=0.04), and the mentor’s graduation record (p=0.010) (**Figure 5C**). There were no differences in respondents of different ranks.

### Factors that PIs believe to be important for obtaining faculty positions

We next asked PIs their opinion on the importance of certain factors for obtaining faculty positions (**Figure 6A**). The majority (13/14) of items queried were determined to be important by more than two-thirds of respondents (**Figure 6A**). Five factors stood out as particularly important, with 90% or more of respondents in agreement: resilience (90% agree, 3% disagree); publications during postdoctoral experience (96% agree, 2% disagree); project idea (97% agree, 1% disagree); hard work (98% agree, 0% disagree); and scientific vision (98% agree, 1% disagree). Amongst the factors that were deemed less important were awards (46% agree, 22% disagree), luck (69% agree, 14% disagree), publications in graduate school (68% agree, 10% disagree), and academic record (75% agree, 11% disagree) (**Figure 6A**).

**Figure 6:**
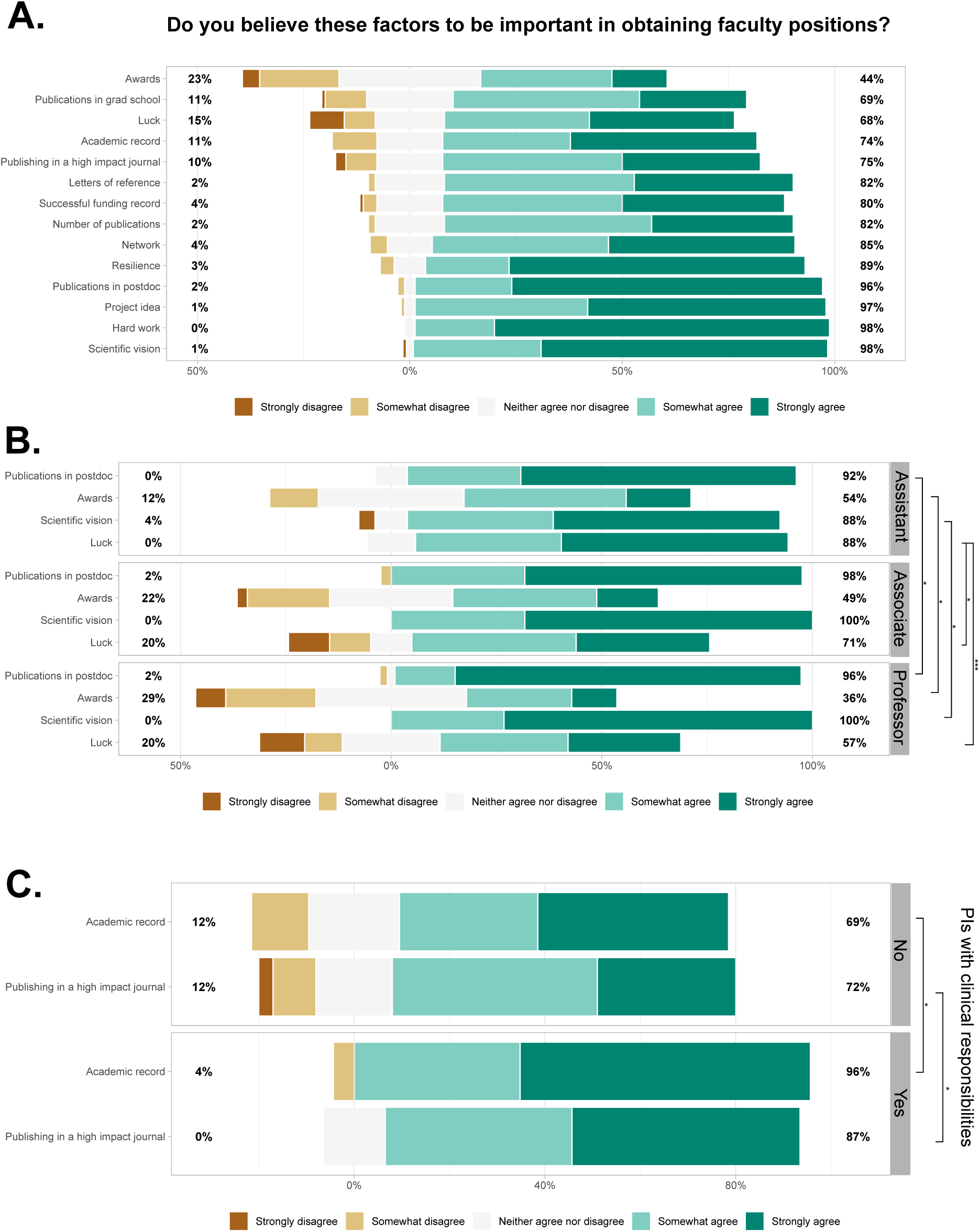
Factors that PIs believed were important in obtaining faculty positions. A. Likert scale chart of queried questions across all PIs. Percentages reflect the aggregate percentage of PIs who “somewhat” or “strongly” agreed or disagreed with an item. B. Likert scale of specific questions whose responses were found to be statistically significant. Results stratified by rank. C. Likert scale of specific questions whose responses were found to be statistically significant. Results stratified by PIs with or without clinical responsibilities. Wilcoxon rank sum test with continuity correction were used to compare conditions (ns represents p > 0.05, * p <0.05, ** p < 0.01, *** p < 0.001).

These responses were not dependent on gender or minority status (**Supplemental File 1**). The responses were generally congruent between professors of different rank. Notably, however, assistant professors were statistically more likely to agree that luck plays a role in obtaining faculty positions (89% agree, 0% disagree) than associate professors (71% agree, 18% disagree, p=0.018) or full professors (58% agree, 18% disagree, p=0.00055) (**Figure 6B**). PIs with clinical responsibilities were significantly more likely to believe that their academic record was important (p=0.019) and publishing in a high impact journal was important (p=0.042) (**Figure 6C**).

### Rating general skills PIs believe contribute to their success

To provide guidance to future PIs about how best to tailor their training, we asked respondents to rank the importance of research-related skills in their success. Amongst these traits were mentorship experience, leadership experience, lab experience, grant-writing experience, networking skills, interviewing skills, negotiating skills, and teaching experience (**Table 2**). PIs ranked grant-writing experience the highest (mean 4.23, 95% CI 4.053 to 4.407) followed by lab experience (mean 3.98, 95% CI 3.76 to 4.19) (**Table 2**). The lowest ranked skills were negotiating skills (mean 2.59, 95% CI 2.35 to 2.83) and teaching experience (mean 1.64, 95% CI 1.42 to 1.86).

**Table 2:** Important PI skills. Table depicting the total number of PI respondents (N between 128-132) and how they rated the importance of certain skills in becoming a PI (from a scale of 1-5). Mean and 95% confidence interval shown.

| **Table 2: Important PI skills** |  |  |  |
| --- | --- | --- | --- |
| Category | Responses (N) | Mean (1-5) | CI (95%) |
| Grant-writing experience | 132 | 4.235 | 0.170 |
| Lab experience | 132 | 3.985 | 0.201 |
| Networking skills | 132 | 3.667 | 0.200 |
| Mentorship experience | 131 | 3.588 | 0.192 |
| Leadership experience | 131 | 3.397 | 0.204 |
| Interviewing skills | 130 | 3.008 | 0.224 |
| Negotiating skills | 130 | 2.631 | 0.230 |
| Teaching experience | 128 | 1.672 | 0.217 |

### Funding is fundamental to becoming a PI

Respondents provided information about the relative contributions of different funding sources that supported their research programs. We found that the average lab is funded mainly by an R01 equivalent (60.0% of funding, 95% CI 54.97% to 64.94%) (**Figure 7A**). PIs also rely on smaller (14.23%, 95% CI 11.29% to 17.17%) and “other” grants (10.43%, 95% CI 7.15% to 13.72%), which account for approximately 26% of an average lab’s funding (**Figure 7A**). However, assistant professors are likely to have smaller research teams. It therefore might be of interest to stratify the responses by academic rank. By doing so, we see that labs from PIs of all ranks rely on ~2 R01 equivalent grants (on average) to run their labs (**Figure 7B**). In addition, we observe that assistant professors rely on statistically significantly fewer R01s to run their lab than associate professors (p=0.025) and full professors (p=0.013) (**Figure 7B, C**). There is no statistically observable difference between funding in labs run by associate professors or full professors (**Figure 7C**). We also found that assistant professors were significantly more likely to rely on their start up packages and smaller grants (**Figure 7C**). The sources of funding were not dependent on gender or minority status (**Figure S3A, B**). However, R01-equivalent grants comprised significantly less of the overall funding for PIs with clinical responsibilities (**Figure S3C**).

**Figure 7:**
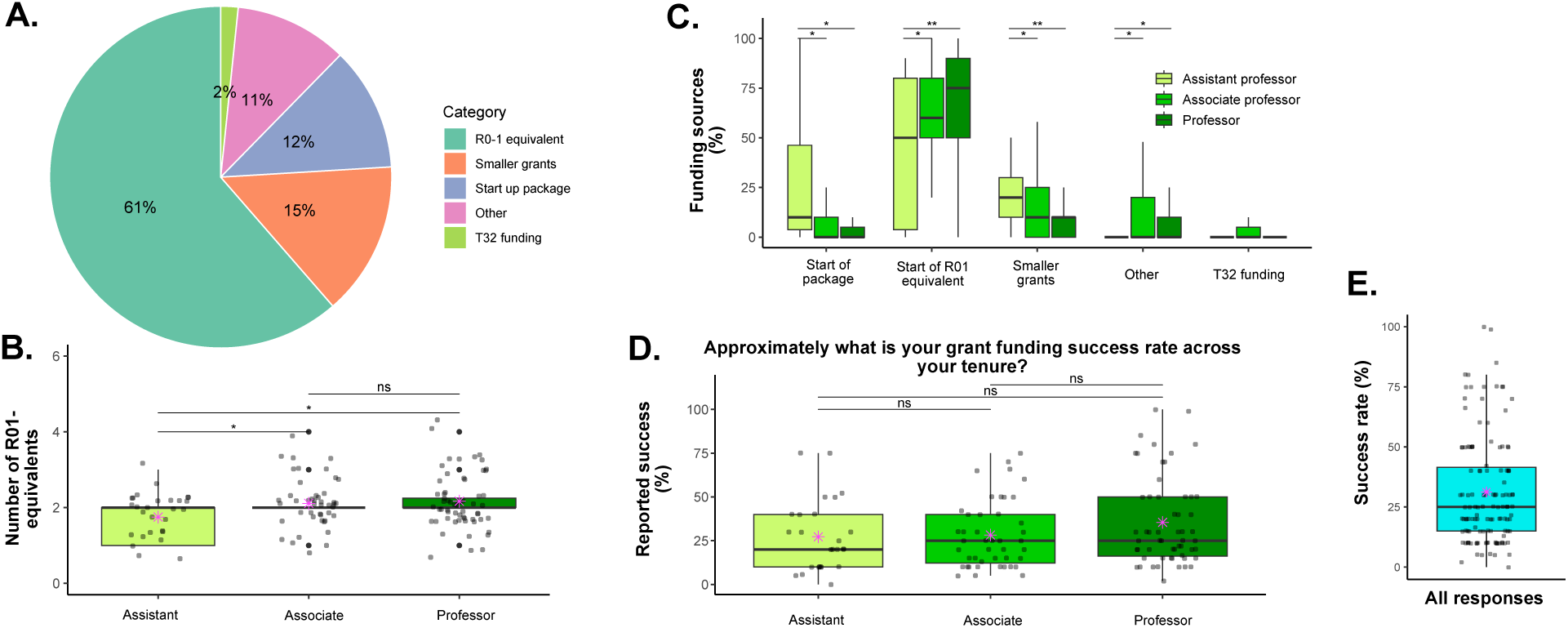
Breakdown of funding sources for PIs. A. A pie graph depicting the funding sources (as a percentage) the average PI uses to cover laboratory operations. B. Boxplots depicting the reported number of R01s needed to fund the labs of PIs. Note: responses are quantized (1, 2, etc), but black dots are distributed for visualization. Mean shown in magenta. Results stratified by academic rank. C. Boxplots depicting the funding sources (as a percentage) PIs use to cover laboratory operations. Results stratified by academic rank. D. Boxplots illustrating the reported grant-funding success (as a percentage). Results stratified by academic rank. Mean shown in magenta. E. Boxplot illustrating the reported grant-funding success for the average PI respondent. Mean shown in magenta. Student T-tests with BH correction were used to compare conditions (ns represents p > 0.05, * p <0.05, ** p < 0.01, *** p < 0.001).

We next asked PIs to reflect on their grant funding success as a PI. It is sometimes assumed that full professors are more successful in acquiring grants than junior faculty (Burns et al. 2018), though the NIH reports differently (“NIH Data Book - Funding Rates”). While we find that full professors report a higher estimated success rate (35.46%, 95% CI 28.71% to 42.22%) than associate professors (28.21%, 95% CI 22.23% to 34.07%) and assistant professors (27.21%, 95% CI 18.94% to 35.48%), the differences were not statistically significant (**Figure 7D**).

We next queried whether demographic factors correlated with different responses about funding success. We found that the funding rate did not differ with minority status (**Figure S3D**). PIs with clinical responsibilities also reported a lower success rate despite having the same proportion of faculty rank (**Figure S3F**).

The average PI at CU Anschutz reported a grant-funding success rate of 30.56% (95% CI to 26.45 to 34.67%) (**Figure 7E**). This self-reported success rate is higher than the NIH average for R01s (“NIH Data Book - Funding Rates” 2024). Surprisingly, women researchers reported higher grant funding success rate than male counterparts (**Figure S3E**), which stands in contrast to published data reporting from NIH, though our data is limited to one institution (“NIH Data Book - Data by Gender” 2024) versus our data limited to one institution.

## DISCUSSION

### Purpose and impact of this study

Traditionally, PhD candidates from the biomedical and basic sciences rely on anecdotal data, the experience of direct and indirect mentors, and limited career-related workshops in order to come to a conclusion about whether or not to become a PI. Previous research has suggested that the more exposure trainees receive about faculty experiences, the more career-prepared they will be (Layton et al. 2020). The goal of this study was to provide more insights about the career paths of PIs conducting work in the biomedical/basic sciences at an R1 stand-alone academic medical institution. Aggregating completed survey responses from 134 PIs from the University of Colorado Anschutz Medical campus, we hoped to have more clearly delineated skills that would be advantageous for future PIs as well as what the daily life of a PI is to help students make an informed decision on whether or not they want to pursue the career of an academic PI in the biomedical/basic sciences.

Overall, we were astonished by the number of PIs that responded, completing a voluntary ten-minute survey with 26 items. We believe the analyses provide a strong foundation for which graduate students can consider the career choice of a PI.

Aside from informing current graduate students considering a PI career, these results may inform best practices for support of trainees/scholars in graduate programs and postdoctoral support groups. Our survey identified that graduate programs and postdoctoral support may benefit from providing more resources and opportunities tailored to writing/obtaining grants, as well as providing the intellectual freedom to develop and explore an independent scientific vision (both of which were rated important for obtaining faculty positions).

It became apparent during our analyses that the study results could also be of interest to institutions seeking methods to help promote/retain faculty at an R1 institution. While we found that some items had responses that were strongly in agreement, significant differences were identified in the study. Many significant differences were noted in responses reflected by academic rank. These data suggest that the significant hardships may be inherent to institutional or career barriers.

We found that junior professors (assistant/associate) were more likely to regret their career choice, less likely to feel financially secure, less likely to recommend becoming a PI, and experience more doubt overall. While the most likely explanation from our data is that academic rank is positively correlated to R01 funding, these attitudes were not eradicated by increased funding at the associate level. These point to other factors that we believe lie beyond the strengths of our survey. Nevertheless, these results may therefore be useful for institutional leaders to identify sources of stress and strategize efforts to improve PI overall wellbeing at early/mid stages of their faculty appointments. Such strategies may involve vertical-integration mentoring and more opportunities or career development for associate professors.

Though the survey was not designed to address physician scientist PIs, the results of this study found that physician scientists may have unique struggles. Nearly 17% of our respondents identified as having clinical responsibilities. Our survey did not specify between MD or MD/PhD physician scientists. Our survey identified that this subset of PIs were less likely to feel financially secure and reported a lower success rate of grant funding (with a smaller proportion reliant on R01 funding). Lower funding rates are a growing trend amongst physician scientists, a vulnerable scientific group (Kwan and Gross 2023). The most likely reason identified in our survey was that they spent significantly less time performing PI duties. However, there are numerous other reasons not assessed in our survey (including administrative support, number of research projects, demands on clinical responsibilities, size of lab, etc). Further studies may uncover the true reason behind these observed differences. Caution should therefore be used to extrapolate this data for physician scientists.

### Limitations of this study

The congruent attitudes as a whole may suggest that there are unifying traits necessary to be a PI, thus lending credibility that advice may not differ significantly from most PIs. Alternatively, given that the data is compiled from a single institution (the University of Colorado Anschutz Medical Campus), these strongly agreed upon factors could be a result of the specific hiring practices or culture at CU Anschutz. For example, in our survey, it was noted that faculty who identified as female were more successful in acquiring grants than their male colleagues, which is not reflected at a national level (Hechtman et al. 2018; van der Lee and Ellemers 2015; “NIH Data Book - Data by Gender” 2024). This could be a direct result of CU School of Medicine’s active efforts to support women researchers, for example, through the Women’s Leadership Training program for women assistant and associate professors and other initiatives such as annual full support for women faculty to participate in the Executive Leadership in Academic Medicine (ELAM) program (resulting in more than 50% of departments led by women chairs). The University of Colorado also strongly values and supports diversity of ideas and identities. This may explain why on most items, there were no significant differences stratified by gender/minority status. However, caution should be used extrapolating gender/minority-specific conclusions from this survey to other institutions without similar cultural values.

It is likely that funding sources, institutional environment, tenureship, or teaching requirements may also vary across institutions. These data would therefore benefit from similar survey comparisons to sister R1 academic institutions as well as non-R1 institutions.

Finally, responses from trainees who chose not to become a PI at an academic institution, became a PI in a non-academic institution, or retired from academia would be useful comparisons. Unfortunately, these populations were not surveyed, as they were out of the scope of this particular study. However, for institutions to improve, it will be important to conduct exit surveys for those who have chosen to leave academia.

### Timing of responses

This survey was conducted for two months (April-June) in 2024. For context, the total NIH budget at this time was $46.1 billion, stable from recent years (Jocelyn Kaiser 2025; “Budget | National Institutes of Health (NIH)” 2025). Our survey results therefore reflect the view of respondents in the context of a stable, healthy NIH funding level. The proposed NIH budget for 2026 is likely to decrease, with proposals of a $27.9 billion allocation. Therefore, our survey results do not explore the impact of decreased NIH budget cuts on how PIs view career choice. Future surveys and comparisons to the present will assess how changes to research funding impact PIs career satisfaction. Nonetheless, the present results inform about factors that PIs consider as important for their success and may provide guidance to current trainees.

## ACKNOWLEDGEMENTS

We would like to thank the biomedical/basic science PIs who helped to develop this survey. They are as follows: Arthur Gutierrez-Hartmann, David Schwartz, Sujatha Jagannathan, Michael McMurray, Jim Costello, Jay Hesselberth, and Julia Cooper. We would also like to thank all the PIs who responded to this survey and took the time to fill it out to help future trainees.

## SUPPLEMENTAL FILES

**Supplemental file 1:** An excel table showing the corrected p values from comparisons made in figures 2, 5, and 6. Questions surveyed are shown in the first column. Comparisons are shown in second row.

**Supplemental file 2:** An excel table of the responses from PIs to the questions: “What is the one reason you chose to become a PI in an academic institution?” (column 1), “What is the hardest thing about being a PI?” (column 2), and “Do you have any additional comments?” (column 3). All PIs agreed to publish these responses.

## SUPPLEMENTAL FIGURES

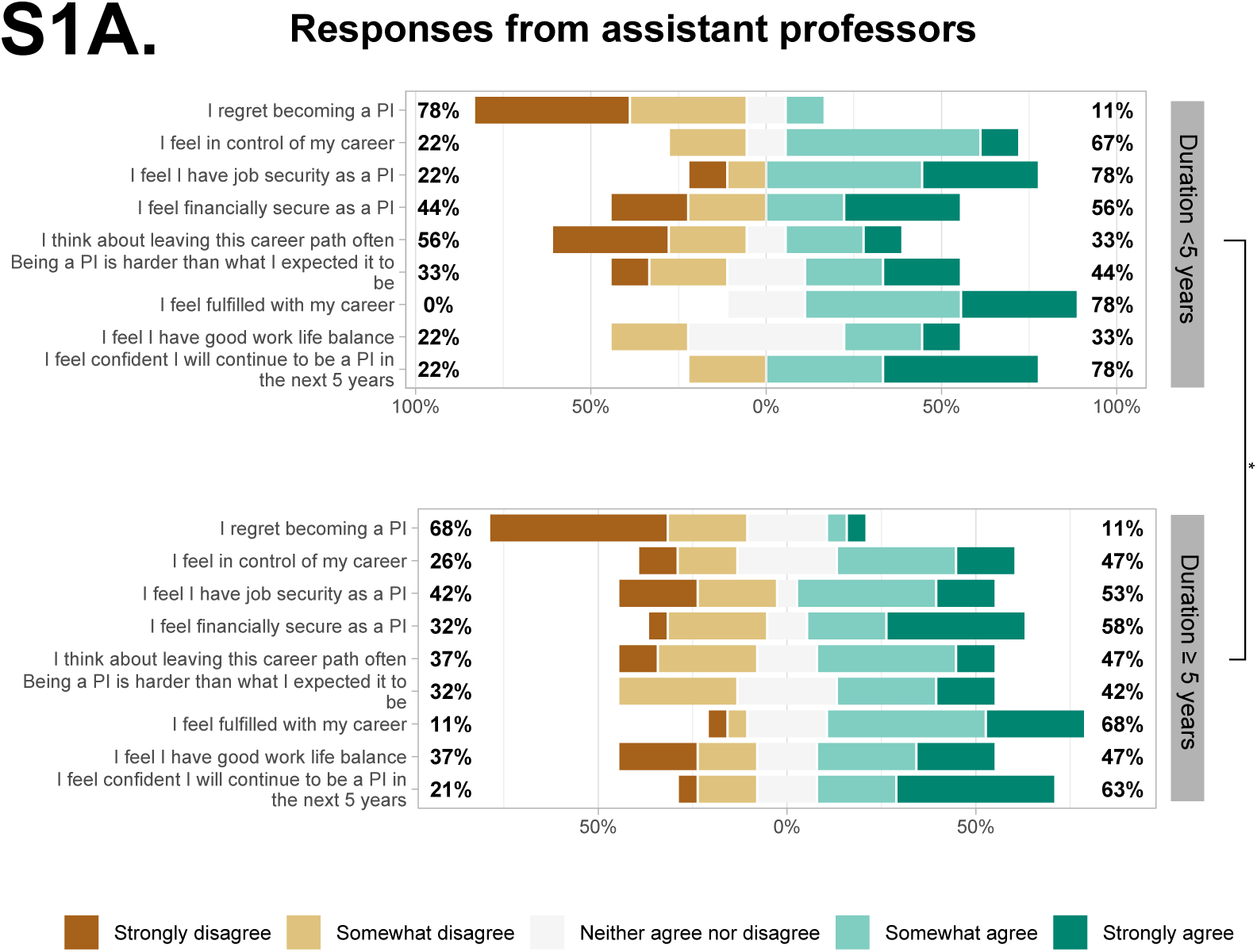

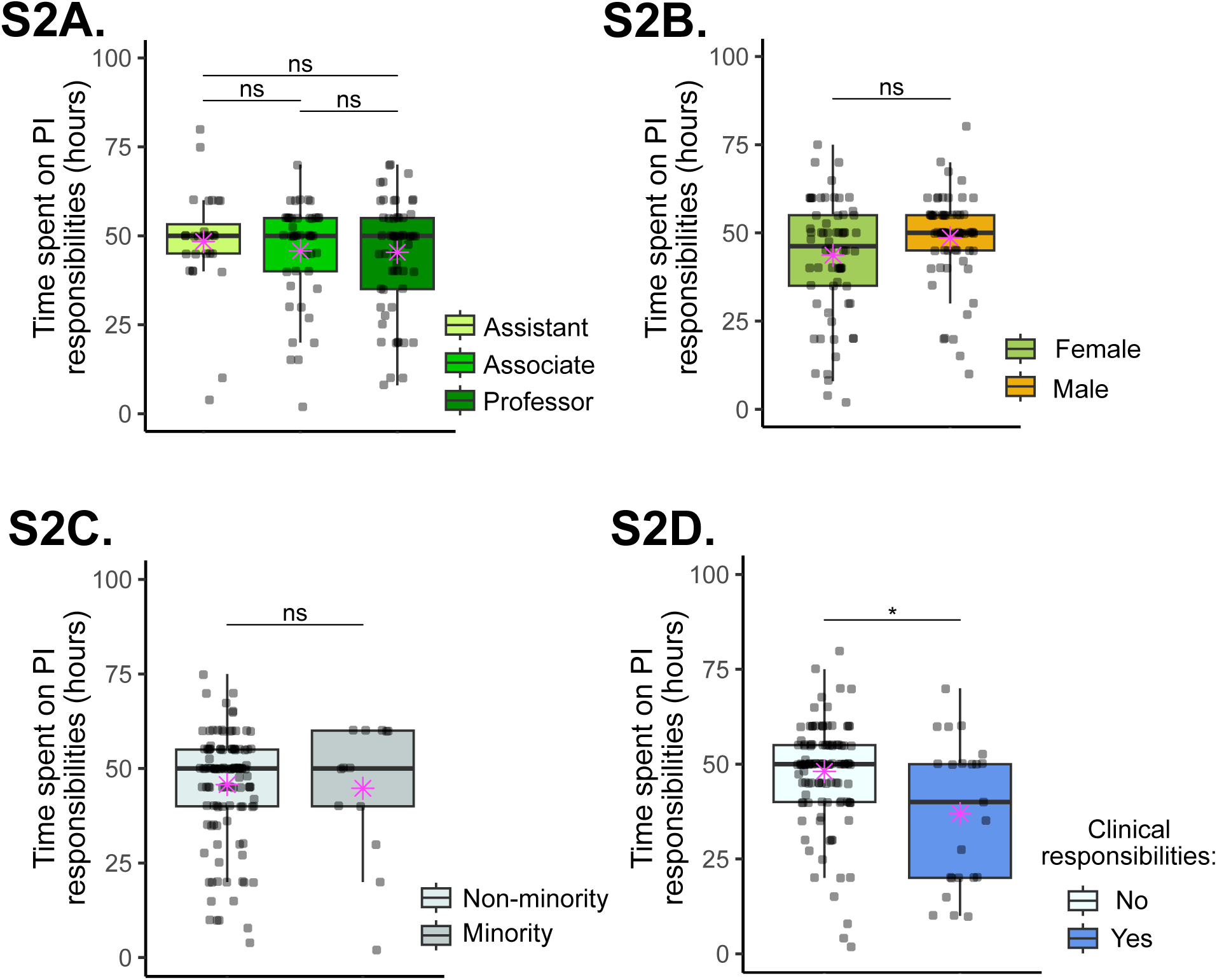
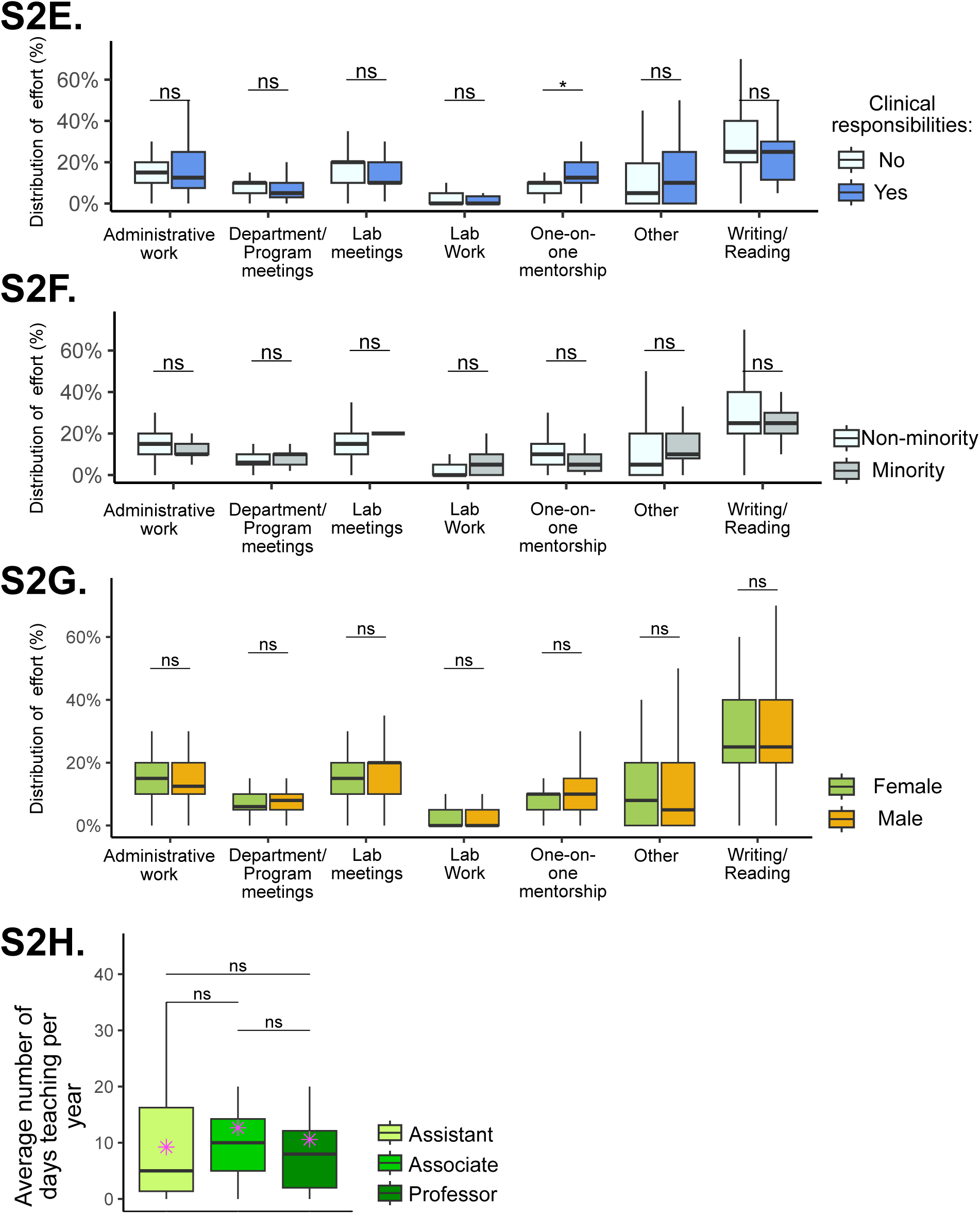

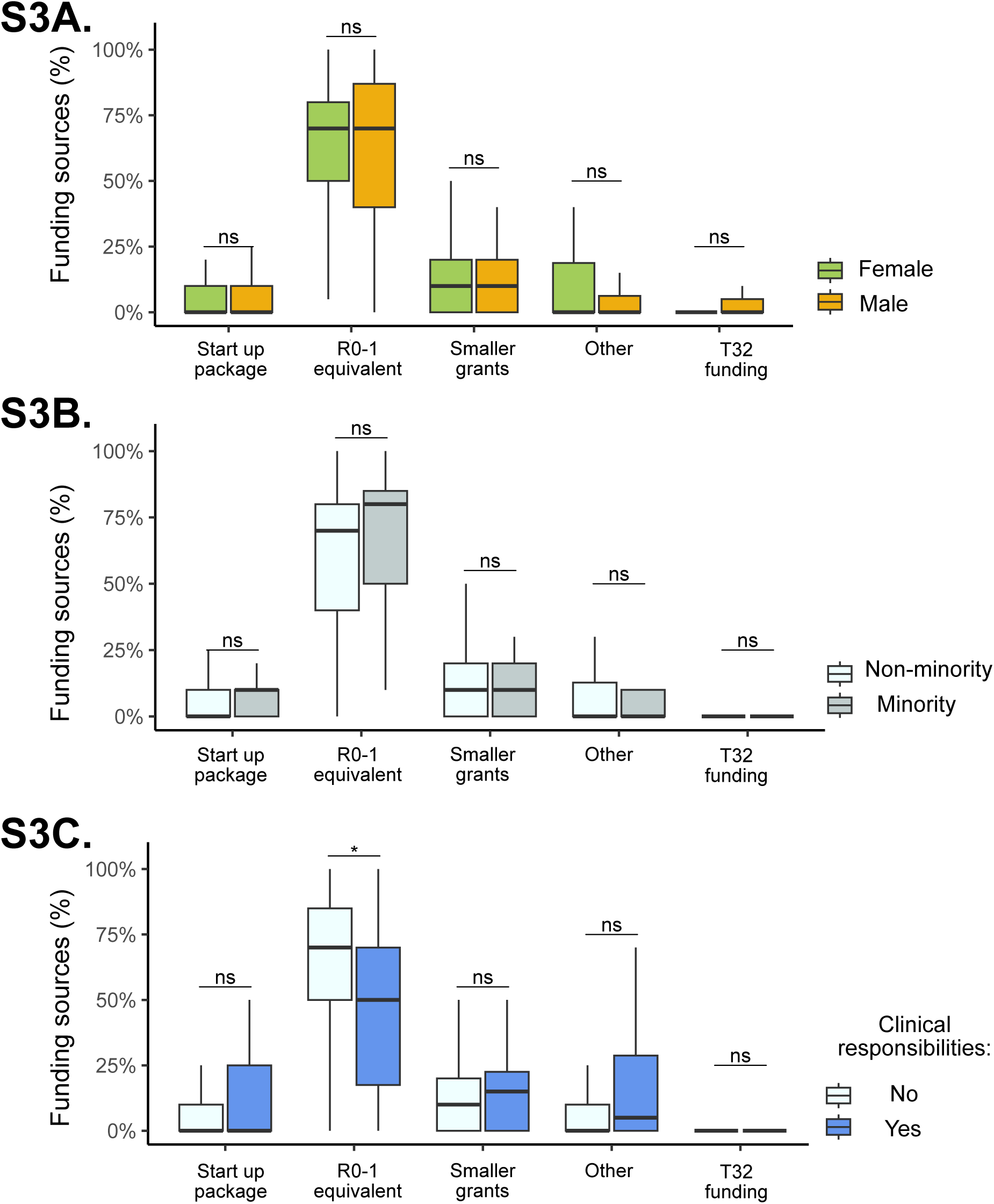
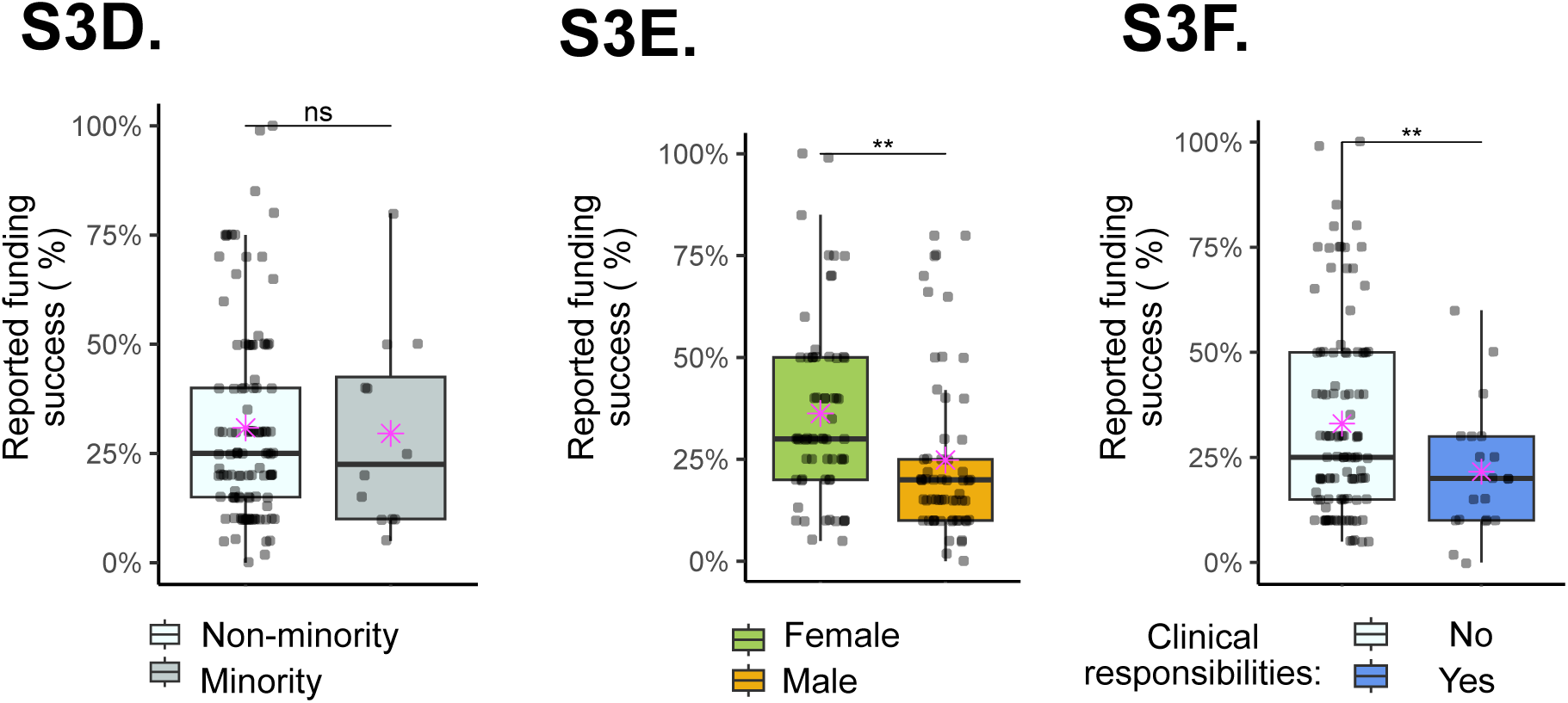

